# A marker-free co-selection strategy for high efficiency human genome engineering

**DOI:** 10.1101/116251

**Authors:** Daniel Agudelo, Lusiné Bozoyan, Alexis Duringer, Caroline C. Huard, Sophie Carter, Jeremy Loehr, Dafni Synodinou, Mathieu Drouin, Jayme Salsman, Graham Dellaire, Josée Laganière, Yannick Doyon

**Affiliations:** Centre Hospitalier Universitaire de Québec Research Center and Faculty of Medicine, Laval University, Quebec City, QC G1V 4G2, Canada.; Research and Development, Héma-Québec, Quebec City, QC G1V 5C3, Canada.; Department of Pathology, Dalhousie University, Halifax, Nova Scotia, B3H 4R2, Canada

## Abstract

Targeted genome editing using engineered nucleases facilitates the creation of *bona fide* cellular models for biological research and may be applied to human cell-based therapies. Broadly applicable and versatile methods for increasing the levels of gene editing in cell populations remain highly desirable due to the variable efficiency between distinct genomic loci and cell types. Harnessing the multiplexing capabilities of CRISPR-Cas9 and Cpf1 systems, we designed a simple and robust co-selection strategy for enriching cells harboring either nuclease-driven non-homologous end joining (NHEJ) or homology-directed repair (HDR) events. Selection for dominant alleles of the endogenous sodium-potassium pump (Na^+^,K^+^-ATPase) that render cells resistant to ouabain is used to enrich for custom modifications at another unlinked locus of interest, effectively increasing the recovery of engineered cells. The process was readily adaptable to transformed and primary cells, including hematopoietic stem and progenitor cells (HSPCs). The use of universal CRISPR reagents and a commercially available small molecule inhibitor streamlines the incorporation of marker-free genetic changes in human cells.

## INTRODUCTION

Designer nucleases such as zinc-finger nucleases (ZFNs), TAL effector nucleases (TALENs), and CRISPR systems greatly simplify the generation of cell lines with tailored modifications enabling the study of endogenous genetic elements in human cells in an unparalleled manner^1-3^. In particular, CRISPR-based technologies enable researchers to introduce double-strand breaks (DSBs) to virtually any DNA sequences using sequence-specific guide RNAs that target Cas9 and Cpf1 nucleases to cleave the matching sequence. The ensuing DSBs activate two competing repair pathways, namely NHEJ and HDR to facilitate the precise modification of the targeted endogenous locus. NHEJ repairs the lesion by directly rejoining the two DSB ends without the need for a repair template resulting in either precise religation or formation of small insertion or deletion mutations (indels) at the break site. In contrast, definite sequence changes can be introduced when the HDR machinery uses an exogenous DNA template with sequence homology to the DSB site to repair the lesion (reviewed in^4^).

While reprograming the CRISPR apparatus requires trivial molecular biology skills, the cellular activity of different guide RNAs can vary considerably, even when targeting adjacent sequences. Computational guide RNA selection is complex and further optimization is required before accurate prediction of the optimal sequences can be achieved, a limitation that is exacerbated when the target site is restricted to a narrow genomic region^5, 6^. Furthermore, the efficiency of delivery and expression of CRISPR components and donor templates can be highly variable between cells of different origin, greatly affecting overall efficacy. Cell type, cell state, and cell cycle stage also play a major role in determining the efficiency of genome editing as human cells dictate repair outcome and have a propensity to favor NHEJ over HDR. Thus, identifying active guide RNAs, screening and isolating clones with the desired genetic modification can be time consuming and costly. In addition, achieving high levels of gene editing is required for accurate phenotypic analysis when studying bulk populations of cells.

At its most basic level, higher genome editing frequencies are associated with higher nuclease levels and activity. Consequently, several approaches have been implemented to capture and isolate these subpopulations of cells in order to expedite the generation of isogenic knockout and knock-in clones. First, co-transfection of fluorescent proteins combined with fluorescence-activated cell sorting (FACS) can be used to isolate nuclease-expressing cells^7^. Second, direct coupling of expression of fluorescent proteins and nucleases via 2A peptide sequences allows for efficient isolation of cell populations with increasingly higher nuclease expression levels, which translates into increasingly higher genome editing rates^8^. Fluorescence-based surrogate target gene reporters have also been successfully used to enrich for cells with high nuclease activity^9, 10^. A limitation of these methods is that single-cell FACS enrichment is not suitable for more sensitive cell lines. In addition, multi-DSBs arising from cleavage of the episomal reporter plasmids are likely to induce a gene-independent antiproliferative response as recently described for CRISPR-Cas9-induced DNA breaks^11^. Finally, enhancing HDR-mediated processes by altering cell cycle parameters, the timing of nuclease expression, chemical inhibition of NHEJ and use of HDR agonists show promising results but their general applicability and absence of negative effects on specificity and genome integrity need to be thoroughly evaluated^12-17^.

Conceptually distinct and elegant genetic approaches based on the creation of “classical” gain-of-function alleles have been developed in the worm *Caenorhabditis elegans.* Termed “co-conversion”/ “co-CRISPR”, these methods increase markedly the odds of detecting a phenotypically silent targeted mutation through the simultaneous “co-conversion” of a mutation in an unrelated target that causes a visible phenotype^18, 19^. It was found that simultaneous introduction of sgRNAs to two different endogenous loci results in double editing events that are not statistically independent. For both NHEJ and HDR-driven modifications, the occurrence of a cleavage and repair event in one locus enhances the probability of finding a heritable mutagenetic event in a second locus in the same animal. A related approach has been described to isolate human cells harboring NHEJ-driven mutations by co-targeting the X-linked hypoxanthine phosphoribosyl-transferase *(HPRT1)* gene^20^. Although relatively efficient, this technique uses a mutagenic chemotherapy drug, does not select for HDR-based events, and may affect the salvage pathway of purines from degraded DNA. Hereditary HPRT1 deficiency is also the cause of Lesch-Nyhan syndrome. More recently, it was found that, in mouse embryonic stem cells, the CRISPR-Cas9-mediated insertion of a drug-selectable marker at one control site frequently coincided with an insertion at an unlinked, and independently targeted site^21^. Still, the insertion of a heterologous selection cassette into the genome of the edited cells may limit applications of this technique. Hence, further refinement of these approaches is needed.

Here we devised a robust marker-free co-selection strategy for CRISPR-driven NHEJ- and HDR-based editing events in human cells by generating dominant cellular resistance to ouabain, a highly potent plant-derived inhibitor of the Na^+^,K^+^-ATPase^22, 23^. The Na^+^,K^+^-ATPase (also known as the sodium-potassium pump), encoded by the *ATP1A1* gene, is the ion pump responsible for maintenance of the electrochemical gradients of Na^+^ and K^+^ across the plasma membrane of animal cells. It is an essential and ubiquitously expressed enzyme functioning as a heteromeric complex consisting of a large α-subunit (ATP1A1), which is responsible for ATP hydrolysis and ion transport and a β-subunit, acting as a chaperone^22, 23^. Cardiotonic steroids (CTSs), such as ouabain (PubChem CID: 439501), constitute a broad class of specific Na^+^,K^+^-ATPase inhibitors prescribed for congestive heart failure for more than 2 centuries^23^. Many mutagenesis and modeling studies have been carried out on the structure-activity relationships of ouabain clearly defining its mechanism of action and leading to the identification of inhibitor-resistant enzymes^22-25^. Using CRISPR-Cas9 and CRISPR-Cpf1, we generated gain-of-function alleles in *ATP1A1* via either the NHEJ or the HDR DNA repair pathways that were used to co-select, through resistance to ouabain, for mechanistically related editing events at a second locus of interest. This strategy was portable to many guide RNAs, independent of cell type, and is a general solution for facilitating the isolation of genome-edited human cells.

## RESULTS

### Editing the endogenous *ATP1A1* locus via either NHEJ or HDR provokes robust cellular resistance to ouabain

Crystal structures of the sodium-potassium pump with bound ouabain and related CTSs have revealed that the inhibitors are wedged very deeply between transmembrane helices in addition to contacting the extracellular surface^22, 23^. Accordingly, random mutagenesis and overexpression studies have pinpointed residues affecting ouabain sensitivity that are localized throughout the protein with clusters of residues conferring the greatest resistance occurring at the H1-H2 transmembrane and extracellular regions^24^. Most prominently, replacement of the border residues (Q118 and N129) of the first extracellular loop with charged amino acids generates highly resistant enzymes when overexpressed in cells^25^ (**Fig. 1a**). However, it is unknown if deletions in this regions can disrupt ouabain binding while preserving the functionality of ATP1A1. Furthermore, it remains to be demonstrated if modification of the endogenous locus, as opposed to overexpression of the mutant enzyme, can result in cellular resistance to ouabain. Therefore, we tested whether targeting this region using the CRISPR-Cas9 system could induce such a phenotype.

**Figure 1.**
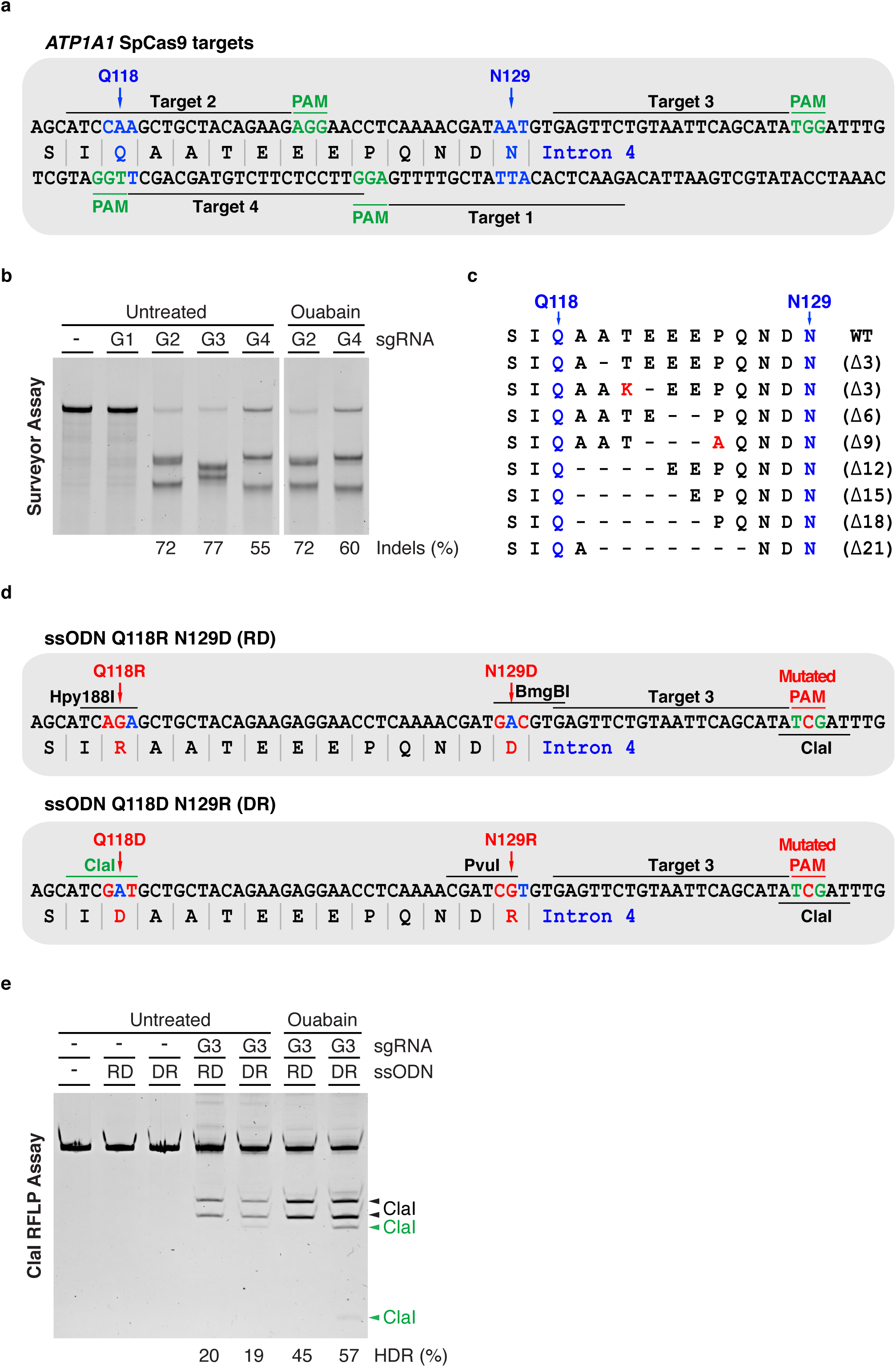
CRISPR-Cas9-driven editing of *ATP1A1* via NHEJ and HDR induces cellular resistance to ouabain. (**a**) Schematic representation of SpCas9 target sites surrounding DNA encoding the first extracellular loop of human *ATP1A1.* Annotated are the positions of residues Q118 and N129, exon/intron boundary, protospacer adjacent motifs (PAM) and four potential SpCas9 target sequences (Targets 1-4). (**b**) The indicated pUC19-based sgRNA expression vectors (G1-G4) (500 ng) were transfected into K562 cells stably expressing wild-type SpCas9 from the *AAVS1* safe harbor locus. Genomic DNA was harvested 10 days post-transfection and the Surveyor assay was used to determine the frequency of SpCas9-induced insertions and deletions, indicated as the *%* Indels at the base of each lane. Where indicated cells were treated with 0.5μM ouabain for 7 days starting 3 days post transfection. An expression vector encoding EGFP (-) was used as a negative control. (**c**) Genomic DNA from (**b**) was amplified by PCR, TOPO cloned and sequenced. (**d**) Schematic representation of the intronic SpCas9 target site G3 and partial sequences of single-stranded oligodeoxynucleotides (ssODNs) donors used to introduce the Q118D/R and N129D/R mutations. Annotated are novel restriction sites introduced to monitor the insertion of ssODN-specified mutations. (**e**) The pUC19-based G3 sgRNA expression vector (500 ng) was co-transfected into K562 cells stably expressing wild-type SpCas9 along with the indicated ssODNs (10 pmole). Cells were treated as in (**b**) and a ClaI RFLP assay was used to determine the frequency of SpCas9-induced HDR at the cleavage site indicated as the % HDR at the base of each lane. An expression vector encoding EGFP (-) was used as a negative control.

We identified two highly active sgRNAs targeting SpCas9 to the exon encoding the first extracellular loop (hereafter named G2 and G4) and one sgRNA targeting the adjacent intron (hereafter named G3) (**Fig. 1a,b**). Starting three days post transfection the cells were treated with ouabain and monitored for survival and growth. Active nucleases targeting the coding sequence (G2 and G4) induced cellular resistance while cells cleaved in the intron with G3 all died within 48 hours (**Fig. 1a,b**). In agreement with previous reports, we have not observed any spontaneous resistance to ouabain treatment^24, 25^. As *ATP1A1* is essential for cell survival, these observations suggest that in frame insertions and deletions were created in the first extracellular loop preventing ouabain binding without crippling enzymatic activity. To assess the spectrum and frequency of targeted mutations generated in these pools of cells we used the TIDE (Tracking of Indels by DEcomposition) method^26^. This analysis revealed that in-frame deletions are selected for over time and upon ouabain treatment and that G2 generates a much more diversified set of mutations than G4, which correlates with a higher fraction of in frame deletions and more robust growth (**Supplementary Fig. 1**). It also shows that the surveyor nuclease assay used to determine the frequency of small insertions and deletions (indels) characteristic of imprecise DSB repair by NHEJ (hereafter, the acronym NHEJ will be used to describe mutagenic repair since the precise religation of the DSBs cannot be detected using this assay) saturates when samples with high levels of modification are tested (**Fig. 1b**). Cloning and sequencing *ATP1A1* alleles from ouabain-resistant cells identified in frame deletion products resulting in disruption of the first extracellular loop of the pump (**Fig. 1c**).

Next, we aimed to test whether it was possible to create gain-of-function alleles by HDR. Reaching a high threshold of HDR in human cells is a major challenge in the genome-editing field since, at the population level, cells favor DSB repair via NHEJ over HDR. Therefore, cleaving within the coding sequence of *ATP1A1* would disfavor the recovery of cells edited through HDR at the expense of cells mutated via NHEJ since ouabain would select for both type of repair events. We took advantage of the highly active sgRNA G3 that targets SpCas9 to the intron to achieve selection exclusively via HDR-driven events (**Fig. 1b,d**). Two single-stranded oligos (ssODNs) were designed to create *ATP1A1* alleles conferring ouabain resistance by replacing both border residues of the first extracellular loop with charged amino acids^25^. The ssODN donors create the double replacements Q118R, N129D (RD) and Q118D, N129R (DR), destroy the protospacer-adjacent motif (PAM) and include additional silent mutations to create restriction sites to facilitate genotyping (**Fig. 1d**). Cas9-expressing cells were co-transfected with sgRNA G3 along with ssODNs and growth was monitored following addition of ouabain. Cells survived and grew robustly only in the presence of the nuclease and either donors. Restriction fragment length polymorphisms (RFLP) assays confirmed the introduction of the desired sequence changes and their enrichment upon ouabain treatment (**Fig. 1e** and **Supplementary Fig. 2**). In addition, increasing the dose of ouabain selected for the double mutants within the population (**Supplementary Fig. 2**). This was not totally unexpected since single mutations at either position confer an intermediate level of resistance^25^. We speculate that it may be possible to select cells having experienced longer gene conversion tracks by increasing the dose of ouabain during selection. Titration of ouabain in the culture medium indicates that cells modified through HDR are resistant to concentration of the drug of at least 1mM, which is more than 100 fold higher than for NHEJ-induced mutations and more than 2000 fold higher than what is needed to kill the cells (0.5 μM). These values correspond to the level of resistance observed in cells overexpressing the mutant enzymes and highlight the wide range of doses that can be used for selection^25^. Thus, optimization of the drug-based selection process for various cell lines is straightforward as it is not necessary to precisely titrate the amount of ouabain required for selection.

**Figure 2.**
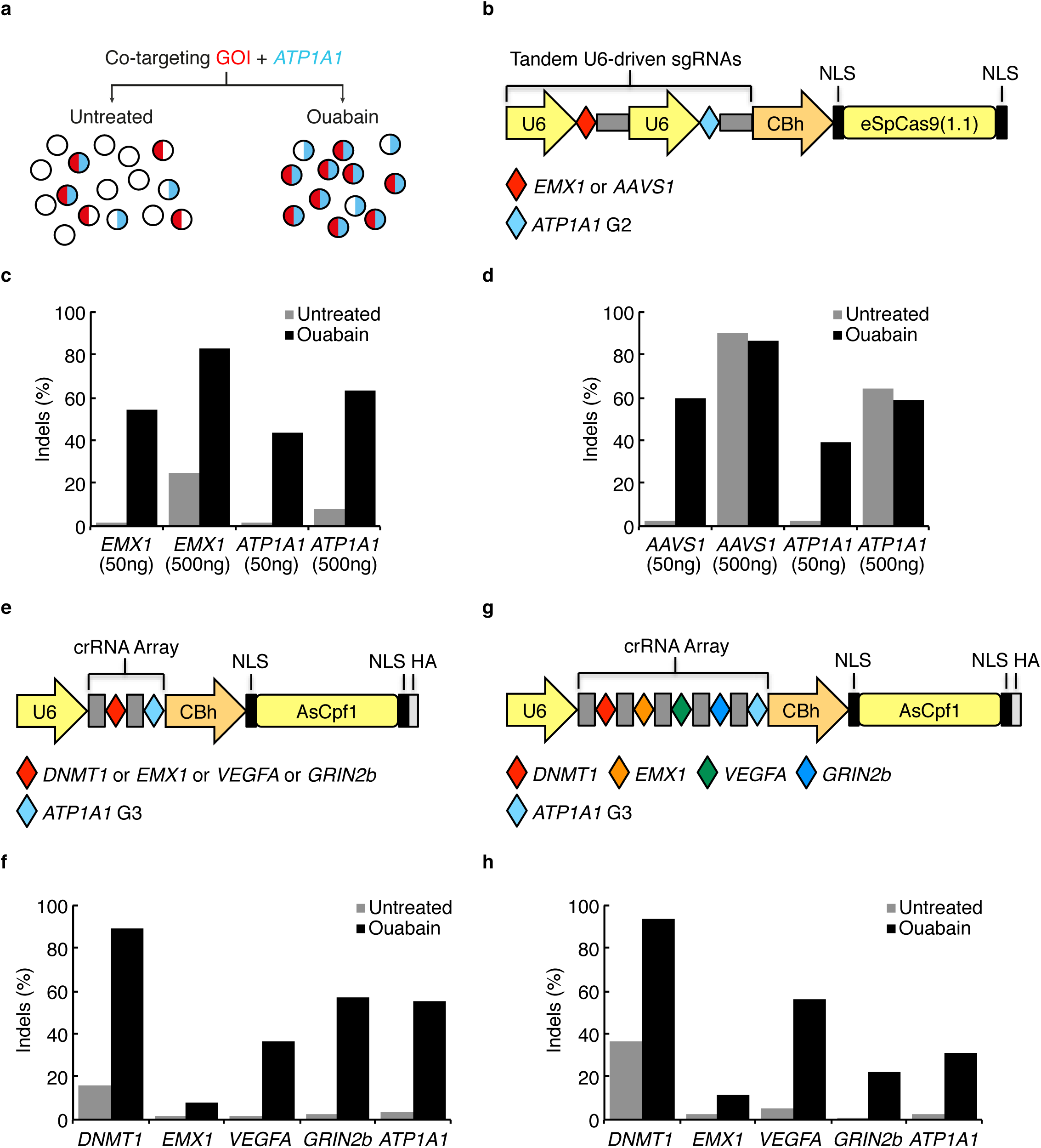
Enrichment of cells with CRISPR-driven targeted mutagenesis by co-editing the *ATP1A1* gene via NHEJ. (**a**) Experimental strategy for the co-enrichment of NHEJ-driven mutations at a second locus. (**b**) Schematic of the dual eSpCas9(1.1) and tandem U6-driven sgRNAs expression vector. (**c**) K562 cells were transfected with 50ng and 500ng of a vector expressing eSpCas9(1.1) and tandem U6-driven sgRNAs targeting *ATP1A1* and *EMX1.* Cells were treated or not with 0.5μM ouabain for 7-10 days starting 3 days post-transfection. Genomic DNA was harvested and the TIDE assay was used to determine the frequency of indels. (**d**) Same as in (**c**) but co-targeting *AAVS1.* (**e**) Schematic of the dual AsCpf1 and U6-driven crRNA array expression vector. (**f**) K562 cells were transfected with 500ng of the vectors shown in (**e**), treated and assayed for indels as (**c**). (**g**) Same as in (**e**). (**h**) Same as in (**f**) but using 1μg of the vector.

These results could be reproduced using the type V CRISPR system from Acidaminococcus sp. Cpf1 (AsCpf1), a single-RNA-guided (crRNA) enzyme that recognizes a TTTN PAM and produces cohesive double-stranded breaks (DSBs)^27^ (**Supplementary Fig. 2**).

In addition, this positive selection was also observed in U2OS, HEK293 and the diploid hTERT-RPE1 cells (**Supplementary Fig. 2** and below). We note that, in selected cells with more than two copies of *ATP1A1,* the fraction of in-frame indels or HDR-alleles can be lower than 50%. For example, in a triploid cell line, the minimal expected signal for these dominant gain-of-function mutations is 33%. Thus, we identified highly active CRISPR-Cas9 and CRISPR-Cpf1 RNA-guided nucleases capable of producing gain-of-function alleles at the *ATP1A1* locus via either NHEJ or HDR.

### Co-selection strategy for enriching cells harboring CRISPR-driven gene disruption events

To test whether selection for the above mentioned gain-of-function alleles in *ATP1A1* can result in co-enrichment of NHEJ-driven mutations at a second locus, we built an all-in-one vector containing tandem U6-driven sgRNA expression cassettes along with CBh-driven high specificity eSpCas9(1.1)^28^ (**Fig. 2a,b**). Cells were transfected with a vector for targeting both *EMX1* and *ATP1A1* and either selected with ouabain or left untreated before genomic DNA was harvested. The frequencies and spectrum of indels in these cell populations was determined by TIDE revealing a marked increase in gene disruption reaching as high as 83% upon selection (**Fig. 2c** and **Supplementary Fig. 3**). In these experiments, transfection of low and high doses of vector was used to simulate the impact of co-selection at a broad range of gene editing frequencies. Similar results were obtained when co-targeting the *AAVS1* and *ATP1A1* loci (**Fig. 2d** and **Supplementary Fig. 3**). In this experiment, the 500ng vector dose transfection was very efficient and starting gene disruption levels were high at both *ATP1A1* and *AAVS1* and no further increase was observed following ouabain treatment (**Fig. 2d**). Surveyor assays performed on all samples corroborated these results (**Supplementary Fig. 4**). Despite reaching almost saturating on-target disruption rates, we could not detect activity at known off-target sites for both *EMX1* and *AAVS1* in these stably modified cell populations^28, 29^ (**Supplementary Fig. 4**). These data indicate that the co-selection process does not negatively impact the enhanced specificity of the eSpCas9(1.1) variant^28^.

**Figure 3.**
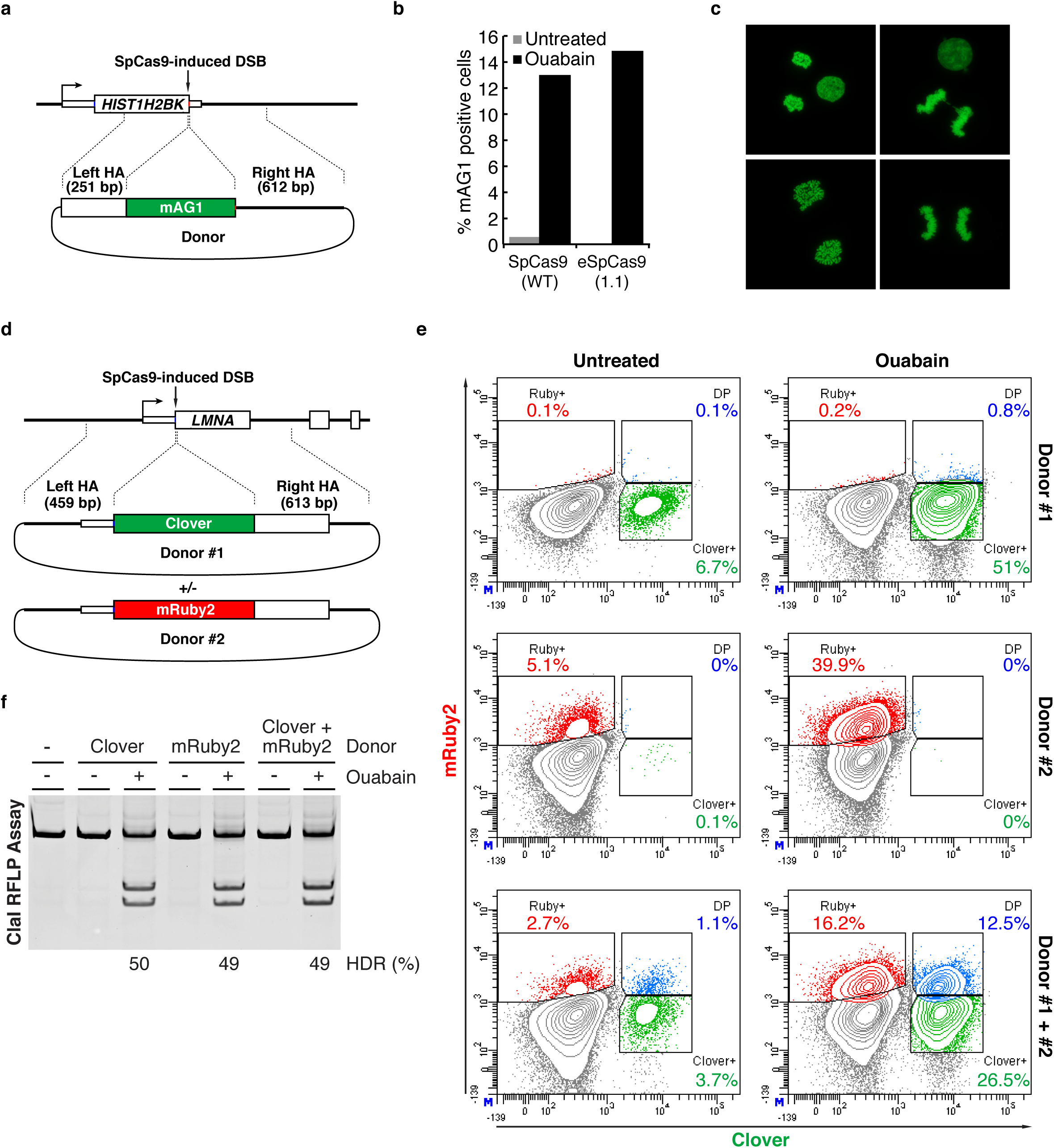
Enrichment of cells with targeted integrations at endogenous loci by co-editing the *ATP1A1* gene via HDR. (**a**) Targeting scheme for the integration of mAG1 to the C-terminus of H2BK. (**b**) K562 cells were transfected with a vector expressing eSpCas9(1.1) and tandem U6-driven sgRNAs targeting *ATP1A1* and *HIST1H2BK* along with *ATP1A1* ssODN RD and a plasmid donor containing a mAG1 cassette. In addition, cells were transfected with a wild-type SpCas9 expression vector and two pUC19-based sgRNA vectors. Cells were treated or not with 0.5μM ouabain for 10 days starting 3 days post-transfection and flow cytometry was used to determine the % of mAG1 positive cells in each population. (**c**) Fluorescence imaging of ouabain treated cells expressing the H2BK-mAG1 fusion. (**d**) Targeting scheme for the integration of Clover and mRuby2 to the N-terminus of lamin. (**e**) K562 cells were transfected with a pX330-based vector (WT SpCas9) expressing the *LMNA* sgRNA, a pUC19-based *ATP1A1* sgRNA vector, along with *ATP1A1* ssODN RD and a plasmid donor containing a Clover or an mRuby2 cassette. Cells were treated as (**b**) and flow cytometry was used to determine the % of Clover and mRuby2 positive cells in each population. (**f**) A ClaI RFLP assay was used to determine the frequency of SpCas9-induced HDR at the *ATP1A1* locus indicated as the % HDR at the base of each lane. Transfection conditions can be found in Supplementary Table 1.

Next, we took advantage of the multiplexing capacity of the CRISPR–Cpf1 system to perform co-selections using all-in-one AsCpf1 expression vectors containing crRNA arrays^30^. We observed improvements in gene disruption efficiency for all four previously published guides, whether expressed in pairs with the *ATP1A1*-targeting guide or as a full array co-expressing five guides simultaneously (**Fig. 2e-h** and **Supplementary Figs. 5** and **6**). These results were reproduced in the HEK293 cell line (**Supplementary Figs. 5** and **6**). As previously observed for SpCas9, the pattern of DNA repair following eSpCas9(1.1) and AsCpf1 cutting at each site is nonrandom, consistent across cell lines and independent of absolute efficacy^31^. AsCpf1 has been shown to be highly specific for its target based on genome-wide specificity assays in human cells so only one known off-target site could be tested amongst the four guides used in the present study^32, 33^. While we observed off-target cleavage for the *DNMT1*-targeting guide in transiently transfected K562 cells we could not detect mutagenesis at this site in ouabain selected cells, even though the on-target activity was superior (**Supplementary Fig. 7**). In contrast, in HEK293, off-target activity was low but apparent (**Supplementary Fig. 7**). Thus, it appears that the co-selection process does not result in overt off-target activity. Collectively, these data show that CRISPR-driven gain-of-function mutations at the endogenous *ATP1A1* gene can be used efficiently for co-selection via NHEJ.

### Robust co-selection for cells harboring CRISPR-driven HDR events

We then tested if selection for cells having experienced a CRISPR-driven HDR event at *ATP1A1* could provide a substantial enrichment for correctly targeted cells at a second locus. We targeted two endogenous genes to generate N- and C-terminal fusions with fluorescent proteins in order to facilitate the quantification of HDR events through FACS-based analysis. To label chromatin, the *HIST1H2BK* locus was targeted to create a C-terminal fusion of H2B with monomeric Azami-Green (mAG1) (**Fig. 3a**). For both wild-type (WT) and eSpCas9(1.1) the fraction of cells expressing the fusion protein increased from below 1% to ~13-15% after ouabain treatment (**Fig. 3b**). The absence of promoter elements in the homology arms of the donor vector along with the clear chromatin-linked fluorescent signal suggests that the process enriched for correctly targeted cells (**Fig. 3c**). Next, we inserted the coding sequence for the green fluorescent protein Clover or the red fluorescent protein mRuby2 after the second codon of the *LMNA* gene, which encodes the lamin A and lamin C isoforms^17^ (**Fig. 3d**). For both Clover and mRuby2 donors we detected a marked increase in signal ranging from ~5-6% to ~40-50% upon ouabain selection and the cells displayed the distinct localization pattern of fluorescence enriched at the nuclear periphery (**Fig. 3e** and **Supplementary Fig. 8**). Co-transfection of the Clover and mRuby2 donors along with the *LMNA* and *ATP1A1*-targeting nucleases permitted to visualize the enrichment of double-positive cells, demonstrating that bi-allelic targeting can be achieved upon ouabain selection (**Fig. 3e**). The level of improvement in gene targeting at the co-selected *LMNA* locus paralleled HDR rates at *ATP1A1* as determined by RFLP assays (**Fig. 3f**). To determine whether the enrichment at the co-selected *LMNA* locus occurs solely for alleles that repaired the DSB via HDR, or whether NHEJ-produced alleles are also enriched for, we performed out-out PCR analysis on ouabain selected samples followed by TOPO cloning and sequencing (**Supplementary Fig. 9**). Sequencing of 44 non-targeted alleles revealed 7 WT sequences and 37 alleles with indels at the predicted cleavage site (**Supplementary Fig. 9**). Thus, NHEJ-produced alleles are also enriched for but a fraction of the cells are likely to have both a targeted and a WT allele. Sequencing of the targeted alleles also indicates that the integration occurred through homologous recombination. Similarly, sequencing of *ATP1A1* alleles revealed 10 WT, 35 NHEJ, and 39 HDR-related events out of 84 reads (**Supplementary Fig. 9**). None of the NHEJ-based deletions extended to the exon. Among HDR events, directional co-conversion of SNPs from the DSB was evident (see **Fig. 1d)**. All 39 clones had incorporated the ClaI site (mutated PAM), 30 had incorporated both the ClaI and the BmgBI sites, and 17 had integrated the 3 RFLP sites. As observed for H2BK, co-selection was efficient for both WT SpCas9 and eSpCas9(1.1) and co-selection using AsCpf1 was also achieved in this system (**Supplementary Fig. 8**). Stimulation of targeted integration of transgene cassettes was also successful at two distinct loci, namely *AAVS1* and *HPRT1* (**Supplementary Fig. 8**). Taken together, these data demonstrate that co-selection for ouabain resistant cells markedly improved the outcome of HDR-driven gene editing experiments, irrespective of the target locus.

### Enabling affinity purification of endogenous protein complexes from co-selected polyclonal cell populations

Having demonstrated that co-selection could generate highly modified cell populations we tested whether functional assays could be performed directly in these pools, thus bypassing single-cell cloning steps. We used tools that we previously developed to build an interface between genome editing and proteomics in order to isolate native protein complexes produced from their natural genomic contexts, an approach that led to the identification of novel interactions in chromatin modifying and DNA repair complexes^34^. Using co-selection, we tagged the enhancer of polycomb homolog 1 (EPC1) and the E1A binding protein p400 (EP400), two essential subunits of the NuA4/TIP60 acetyltransferase complex that promote homologous recombination by regulating 53BP1-dependent repair^35^ (**Fig. 4a,b**). Out-out PCR-based assays and western blotting confirmed the correct integration of the affinity tag at both loci and the enrichment of tagged cells upon ouabain treatment (**Fig. 4c,d**). Nuclear extracts were prepared from the cell pools as well as from WT K562 cells (Mock), and subjected to tandem affinity purification (TAP). EPC1 and EP400 complex subunits separated by SDS-polyacrylamide gel electrophoresis (SDS-PAGE) could be unambiguously identified on silver stained gels and are virtually identical to the patterns obtained from clonal cell lines^34^ (**Fig. 4e**). Successful tagging via co-selection was also achieved in HEK293 cells (**Supplementary Fig. 10**). These results represent an additional step toward high-throughput genome-scale purification of native endogenous protein complexes in human cells.

**Figure 4.**
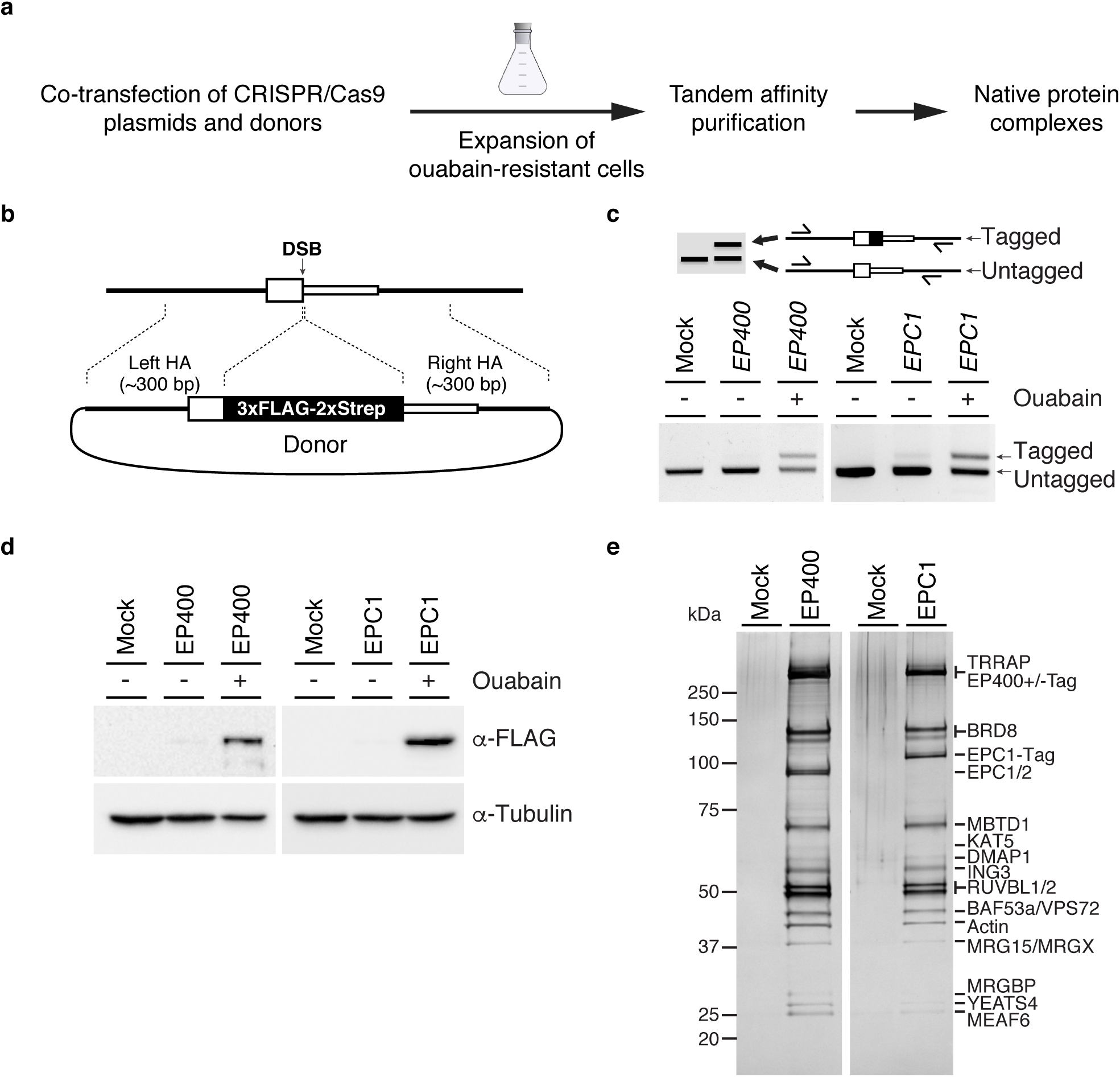
Endogenous tagging and tandem affinity purification of the native NuA4/TIP60 complex from co-selected cell pools. (**a**) Schematic representation of the experiment. (**b**) Gene tagging scheme. (**c**) K562 cells were transfected with a vector expressing eSpCas9(1.1) and a sgRNA targeting *ATP1A1* along with a pUC19-based sgRNA vector for *EPC1* or *EP400* and *ATP1A1* ssODN RD and a plasmid donor containing a 3xFLAG-2xStrep epitope sequence. Cells were treated or not with 0.5μM ouabain for 10 days starting 3 days post-transfection and a PCR-based assay (out-out PCR) was used to detect targeted integration (TI) of the tag sequence at the C-terminus of EPC1 and EP400. Primers are located outside of the homology arms and are designed to yield a longer PCR product if the tag is inserted. (**d**) Western blot analysis on whole cell extracts to detect the expression of EPC1-tag and EP400-tag proteins. The FLAG M2 antibody was used to detect tagged proteins and the tubulin antibody was used as a loading control. (**e**) Silver stained SDS-PAGE showing the purified EPC1 and EP400 complexes.

### Efficient enrichment of gene-edited human hematopoietic stem and progenitor cells (HSPCs)

To explore the potential for clinical translation of our method we tested the ouabain selection strategy during *ex vivo* expansion of cord blood-derived human hematopoietic stem and progenitor cells (HSPCs). Genome editing in HSPCs by homologous recombination remains challenging but offers the possibility to treat several genetic diseases^36^. We used previously developed tools to introduce a mutation in the beta-globin (HBB) gene causing sickle cell disease (SCD)^37-39^ (**Fig. 5a**). Purified human CD34^+^ cells were electroporated with preformed SpCas9 ribonucleoprotein complexes (RNPs) containing synthetic crRNAs and tracrRNAs targeting *ATP1A1* and *HBB* and ssODNs and expanded *ex vivo* with or without ouabain. Cells were cultivated in presence of UM171 to promote expansion and maintenance of primitive progenitors during selection^40^. RFLP-based assays clearly indicate that cells edited at the *HBB* locus could be efficiently enriched using ouabain, a phenomenon observed in cells isolated from various donors and using different ssODNs (**Fig. 5b,c** and **Supplementary Fig. 11**). These results support the notion that the process could be adapted for use in preclinical studies. Critically, these data demonstrate that the procedure is applicable to diploid primary cells.

**Figure 5.**
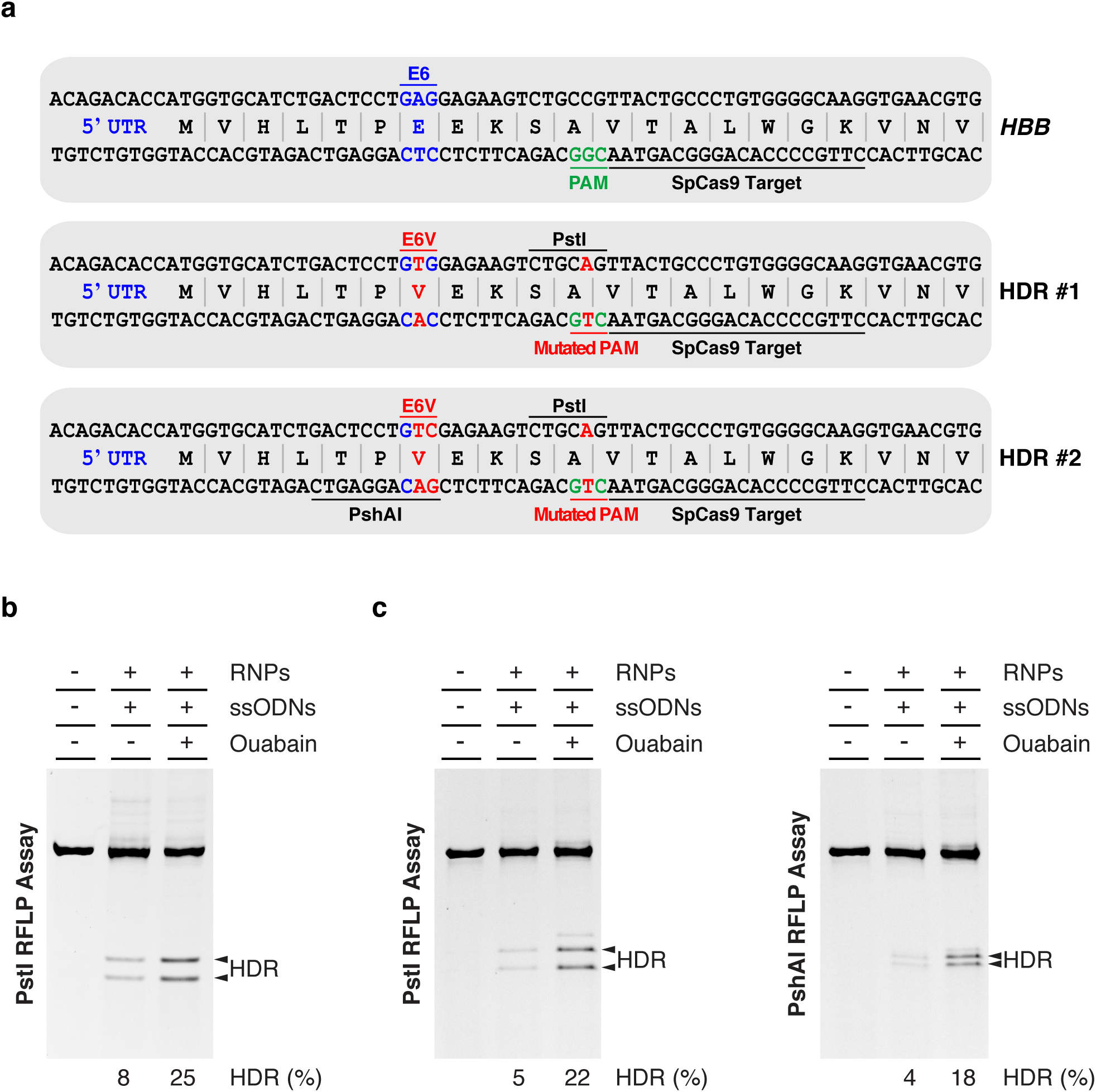
Enrichment of HDR-driven events in primary human cord blood (CB) CD34+ cells upon selection with ouabain. (**a**) Schematic representation of SpCas9 target site in *HBB* and predicted HDR outcomes dictated by two different ssODNs donors used to introduce the E6V mutation. Annotated are the positions of E6 residue, 5’ UTR, protospacer adjacent motifs (PAM) and novel restriction sites introduced to monitor the insertion of ssODN-specified mutations. HDR#2 bears an additional PshAI site to directly monitor conversion of the E6V mutation. (**b**) Cultured CD34+ cells were electroporated with *ATP1A1* and *HBB* RNPs along with *ATP1A1* ssODN RD and *HBB* ssODN#1 and grown in the presence of 0.5μM ouabain starting 5 days post-transfection. Genomic DNA was harvested at 13 days post-transfection and a PstI RFLP assay was used to determine the frequency of SpCas9-induced HDR, which is indicated as the % HDR at the base of each lane. Recombinant Cas9 was used as a negative control (-). (**c**) CD34+ cells were treated as in (**b**) but using *HBB* ssODN#2 and assayed by RFLP for both PstI and PshAI sites.

## DISCUSSION

Here we demonstrate that the creation of gain-of-function alleles at the *ATP1A1* locus with CRISPR-Cas9 and Cpf1 can be robustly selected for using a highly potent therapeutic small molecule in order to enrich for custom modifications at another unlinked locus of interest, substantially increasing the recovery of engineered cells. A defining aspect of our system is that the co-selection process can be initiated through NHEJ- or HDR-driven events independently. One only has to switch between two validated sgRNAs targeting juxtaposed regions of *ATP1A1* and include an ssODN in the transfection reaction to co-select for HDR- instead of NHEJ-driven events. Perhaps more importantly, we show that the co-selection strategy markedly enriches for cells modified via homologous recombination, a clear limitation in the field. This was demonstrated at endogenous loci for five different types of HDR-driven genome editing events; (*i*) RFLP knock-in, *(ii)* gene trapping, *(iii)* targeted integration of an autonomous expression cassette, *(iv)* protein tagging with fluorescent markers, and (*v*) epitope tagging. As shown in the examples above, the extent of enrichment for HDR-edited cells varied at different sites and according to the type of modification created at the locus of interest. Of particular importance, robust selection is achieved without the use of exogenous DNA markers making it potentially compatible with therapeutic applications. Further work and optimization will reveal if the gene editing and selection process is compatible with a variety of primary cell types. The well-defined mechanism of action of ouabain acting on a nonsignaling ion pump independently of proliferation is another distinct advantage of the method. Ouabain treatment kills cells within 48 hours of exposure and targeted cells display no apparent growth delay resulting from the selection process. Accordingly, the point mutations engineered to confer ouabain resistance are naturally occurring in mice, rats, monarch butterflies, leaf beetles, and some toads^41^. In addition, these mutant enzymes function normally, as shown by ^86^Rb^+^ uptake and ATP hydrolysis assays^25^. The turnover of ATP1A1 at the plasma membrane appears to be rapid since ouabain can be added to the culture medium as early as 15 hours post transfection of the CRISPR components, even if they are encoded on plasmids (data not shown). Although not formally tested in this study, the process should be compatible with any engineered nuclease platforms and any cell type since *ATP1A1* is ubiquitously expressed.

These results corroborate earlier observations that cells proficient at completing one genomic manipulation have an increased probability of completing a second, independent genomic manipulation, provided they are mediated by sufficiently similar mechanisms of DNA repair^18, 19, 21^. Powerful co-selection schemes such as the one described in this study may offer the opportunity to efficiently correct a mutation “at distance” and avoid complications caused by on-target NHEJ-based mutagenesis of the non-corrected allele. Since about 80% of human exons are < 200 bp in length, it should be possible to cleave within an intron to induce recombination in the juxtaposed exon. Provided the intronic sequence is not repetitive and indels in the intron do not affect gene expression such a strategy should be considered. The concept of creating and selecting for dominant gain-of-function alleles of endogenous genes and its use to enrich for custom modifications at unlinked loci of interest could be broadened to include other protein/drug or cell surface marker/antibody combinations. For example, one could achieve the co-enrichment of edited cells *in vivo* after chemo-selection of transplanted cells modified to express the P140K variant of human O(6)-methylguanine-DNA-methyltransferase (MGMT)^42^. Technically, it is interesting to note that this residue is encoded next to an intron-exon boundary and could be targeted in a similar fashion as *ATP1A1* to favor enrichment of cells edited via HDR (**Supplementary Fig. 11**).

Remarkably, robust co-selection for NHEJ-based on-target mutagenesis does not yield overt off-target activity when enhanced fidelity variants of SpCas9 are used. In fact, we did not detect mutagenesis at sites where WT SpCas9 is known to cleave. It also appears that the co-enrichment of indels at off-target sites is limited compared to the one observed at on-target loci, as shown for AsCpf1. As the amount of CRISPR components can be titrated to improve the ratio of on-target to off-target mutation rates, an unexpected advantage of the co-selection strategy may be to achieve this balance without reducing the final on-target efficacy. This will hold true if transfecting lower amounts of plasmids results in lower levels of the nuclease on a per-cell basis. Translocations between *ATP1A1* and the locus of interest will occur at some frequency and there is a slight probability that this event will be found in the selected population.

In conclusion, we established a simple method that can be broadly applied to increase the frequency of genome editing events in human cells. It is easy to implement in any laboratory, as it only requires a universal set of CRISPR reagents and an inexpensive commercially available drug. Furthermore, by design, it was built to be compatible with any nuclease platform and most other strategies developed to facilitate the use of this powerful technology.

## METHODS

### Cell culture and transfection

K562 were obtained from the ATCC and maintained at 37 °C under 5% CO_2_ in RPMI medium supplemented with 10% FBS, penicillin-streptomycin and GlutaMAX. U2OS cells were obtained from the ATCC and maintained at 37 °C under 5% CO_2_ in McCoy’s 5A medium supplemented with 10% FBS, penicillin-streptomycin and GlutaMAX. HEK-293 LTV were purchased from Cell Biolabs and hTERT RPE-1 cells were a kind gift from Amélie Fradet-Turcotte. Both cell lines were maintained at 37 °C under 5% CO_2_ in DMEM medium supplemented with 10% FBS, penicillin-streptomycin and GlutaMAX. Cells (2E5 per transfection) were transfected using the Amaxa 4D-Nucleofector (Lonza) per manufacturer’s recommendations. Transfection conditions used in co-selections via HDR can be found in Supplementary Table 1. Ouabain octahydrate (Sigma) was dissolved at 50 mg/ml in hot water and stored at −20 °C. Working dilutions were prepared in water and added directly to the culture medium.

### CRISPR-Cas9 and Cpf1 reagents

The CAG-driven human codon optimized Cas9 nuclease vectors hCas9 was a gift from George Church (Addgene plasmid # 41815)^43^. The expression cassette was transferred to AAVS1_Puro_PGK1_3xFLAG_Twin_Strep (Addgene plasmid *#* 68375)^34^ in order to establish the K562 cell line constitutively expressing Cas9. All sgRNA expression vectors were built in the SP_gRNA_pUC19 backbone (Addgene plasmid # 79892)^34^, with the exception of gRNA_AAVS1-T2 which was a gift from George Church (Addgene plasmid # 41818)^43^. Design of sgRNAs was assisted by the GPP Web Portal (http://portals.broadinstitute.org/gpp/public/) and the MIT web-based CRISPR design tool (http://crispr.mit.edu/). Sequences of the guides are provided in Supplementary Table 2. The DNA sequence for the guides were modified at position 1 to encode a ‘G’ due to the transcription initiation requirement of the human U6 promoter when required. The reagents targeting the *LMNA* gene have been described^17^. To test the high specificity eSpCas9(1.1) variant, guide sequences were cloned into the eSpCas9(1.1)_No_FLAG vector (Addgene plasmid # 79877). The human codon optimized AsCpf1 ORF from the nuclease vector pY010, a gift from Feng Zhang (Addgene plasmid # 69982)^27^, was transferred to AAVS1_Puro_PGK1_3xFLAG_Twin_Strep (Addgene plasmid # 68375)^34^ in order to establish the K562 cell line constitutively expressing Cpf1 from a CAG promoter. A crRNA expression vector was built in the pUC19 backbone by linking the AsCpf1 5’DR to a human U6 promoter^27^. The guide sequences where cloned downsteam of the 5’DR. Guide for AsCpf1 were designed using Benchling (https://benchling.com/academic) and sequences are provided in Supplementary Table 3. The dual AsCpf1 and U6-driven crRNA array expression vector was built in pY036 (a gift from Feng Zhang) as described^30^. Desalted ssODNs (Supplementary Table 4) were synthesized as ultramers (IDT) at 4 nmole scale. All plasmid donor sequences contain short homology arms (<1 kb) and have been modified in order to prevent their cleavage by Cas9 as described^34^. The only exception to this rule was for the targeting of Clover to the *LMNA* locus using AsCpf1. eSpCas9(1.1) and AsCpf1 expression vectors targeting *ATP1A1* have been deposited to Addgene (**Supplementary Fig. 13**).

### Surveyor nuclease, RFLP knock in assays, out-out PCR assays, and TIDE analysis

Genomic DNA from 2.5E5 cells was extracted with 250 μl of QuickExtract DNA extraction solution (Epicentre) per manufacturer’s recommendations. The various loci were amplified by 30 cycles of PCR using the primers described in Supplementary Table 5. Assays were performed with the Surveyor mutation detection kit (Transgenomics) as described^44^. Samples were separated on 10% PAGE gels in TBE buffer. For RFLP assays, the PCR products were purified and digested with the corresponding enzyme and resolved by 10% PAGE. Gels were imaged using a ChemiDoc MP (Bio-Rad) system and quantifications were performed using the Image lab software (Bio-Rad). TIDE analysis was performed using a significance cutoff value for decomposition of p<0.005^26^. To analyze targeted integration via out-out PCR, genomic DNA extracted with QuickExtract DNA extraction solution was subjected to 30 cycles of PCR for *LMNA* and 35 cycles for *EPC1* and *EP400* using the primers described in Supplementary Table 6. Amplicons were loaded on 1% agarose gels in TAE buffer.

### Flow cytometry

The frequency of EGFP, mAG1, Clover, and mRuby2 expressing cells was assessed using a BD LSR II flow cytometer and gated for viable cells using 7-aminoactinomycin D (7-AAD).

### TAP

Nuclear extracts and purifications were performed from ~5E8 cells as described^34^. Approximatively 1/30 of the final eluates were loaded without prior precipitation on Bolt 4–12% Bis-Tris gels (Life Technologies) and ran for 45 minutes at 200 volts in MOPS buffer. Silver staining was performed using the SilverQuest kit (Life Technologies).

### Human cord blood (CB) CD34+ cell collection and processing

Human umbilical CB samples were collected from donors consenting to procedures approved by the Research Ethic Board of the CHU de Québec - Université Laval. Mononuclear cells were first isolated using Ficoll-Paque Plus density centrifugation (GE Healthcare). Human CD34^+^ hematopoietic stem and progenitor cells (HSPCs) were isolated using CD34 positive selection according to manufacturer’s instructions (EasySep^™^, StemCell Technologies). Purified CD34+ cells were cryopreserved in Cryostor CS10 (StemCell Technologies).

### HSPC culture, editing and selection

CD34+ HSPCs were thawed and cultured in StemSpan ACF (StemCell Technologies) supplemented with 100ng/ml SCF (Feldan), 100ng/ml FLT3-L (Peprotech), 50ng/ml TPO (StemCell Technologies), 10μg/ml LDL (StemCell Technologies) and 35nM UM171 (StemCell Technologies) for 16-24h after thawing. Cells were then nucleofected with Cas9 RNP and ssODNs using the Amaxa 4D Nucleofector X unit (Lonza) and the E0-100 program according to manufacturer’s recommendations. Following nucleofection, cells were incubated at 30°C for 16h and then transferred at 37°C for 2 days. From day 3 post-nucleofection, HSC were transferred to StemSpan ACF (StemCell Technologies) containing 1X StemSpan CD34+ Expansion supplement (StemCell Technologies) and UM171. Ouabain (0.5μM) was added to the cells during culture media change at day 5 post-nucleofection, replenished every 3-4 days and maintained for 8 days prior to RFLP analysis at D14 post thaw.

### Synthetic crRNA and tracrRNA complexes and ribonucleoprotein (RNP) formation for editing in HSPCs

The Alt-R CRISPR system (Integrated DNA Technologies) was used to produce the *HBB*^37–39^ (CTTGCCCCACAGGGCAGTAA) and *ATP1A1* G4 (GTTCCTCTTCTGTAGCAGCT) guides. crRNA and tracrRNA were first resuspended to 200μM stock solutions in Nuclease-Free IDTE Buffer (Integrated DNA Technologies). For crRNA:tracrRNA complex formation, the two RNA oligos were mixed in equimolar concentrations, heated at 95°C for 5 minutes and transferred to ice immediately. Immediately prior to nucleofection, 50pmol of Cas9 protein (Integrated DNA Technologies) was incubated with 125pmol of crRNA:tracrRNA complexes at RT for 10 minutes to form the RNP complexes along with 100pmole of each ssODNs. Note that the *ATP1A1* G3 guide was not active in the Alt-R CRISPR system (Integrated DNA Technologies), thus we used the *ATP1A1* G4 guide to make RNPs. This guide generates low levels of ouabain resistance via NHEJ in HSPCs and robust cell growth is only observed in the presence of ssODN donors (Supplementary Table 4).

## ACKNOWLEDGMENTS

This study was supported by grants from the Natural Sciences and Engineering Research Council of Canada (RGPIN-2014-059680) to Y.D. and the Canadian Institutes of Health Research (CIHR; MOP-84260) to G.D. Salary support was provided by the Fonds de la recherche du Québec-Santé (FRQS) for D.A. and Y.D. and by a bridge grant from the Beatrice Hunter Cancer Research Institute (BHCRI) for J.S. We thank Sabine Elowe, Amélie Fradet-Turcotte, Daniel Durocher, Jacques Côté, and Christian Beauséjour for critical reading of the manuscript as well as the staff at the Hôpital St-Francois d’Assise for their precious help in collecting UCB units.

## AUTHOR CONTRIBUTIONS

Conceptualization, Y.D.; Methodology, D.A., L.B., A.D., C.C.H., S.C., J.L., and Y.D.; Investigation, D.A., L.B., A.D., C.C.H., S.C., J.L., D.S., M.D., and Y.D.; Resources, J.S., and G.D.; Writing – Original Draft, Y.D.; Writing – Review and Editing, D.A., L.B., A.D., S.C., J.L., G.D., J.L., and Y.D.; Supervision, J.L., and Y.D.; Funding Acquisition, J.L., and Y.D.

